# BulkECexplorer: a bulk RNAseq compendium of five endothelial subtypes that predicts whether genes are active or leaky

**DOI:** 10.1101/2023.10.26.564202

**Authors:** James T. Brash, Guillermo Diez-Pinel, Alessandro Fantin, Christiana Ruhrberg

## Abstract

Transcriptomic data obtained by single cell (sc) RNAseq or bulk RNAseq can be mined to understand the molecular activity of cell types. Yet, lowly expressed but functional genes may remain undetected in RNAseq experiments for technical reasons, such as insufficient read depth or gene drop out in scRNAseq assays. By contrast, bulk RNAseq assays may detect lowly expressed mRNA transcripts thought to be the biologically irrelevant products of leaky transcription. To more accurately represent a cell’s functional transcriptome, we propose compiling many bulk RNAseq datasets into a compendium and applying established classification models to predict whether the detected genes are likely active or leaky in that cell type. Here, we have created such a compendium for vascular endothelial cells from several mouse and human organs, termed the BulkECexplorer.

## Introduction

RNA sequencing (RNAseq) has emerged as the leading method to interrogate transcriptomic signatures for specific physiological and pathological states of cell populations. Single cell (sc) RNAseq compendia provide useful resources to identify cell types via their transcriptomic signature and to compare the expression pattern of a gene at single cell level across a range of cell types and tissues (e.g., Griffiths et al., 2018). By contrast, bulk RNAseq has been widely used to determine average transcript levels in a given cell population, for example to compare different genes within one cell type or transcriptomic changes after experimental manipulation (e.g., Kulahoglu and Brautigam, 2014). A wealth of vascular endothelial cell (EC) transcriptome data has been generated using RNAseq (e.g., Khan et al., 2019, He et al., 2018, Jambusaria et al., 2020). Much of these data are publicly available and can be mined to generate new insights into EC biology. Considering the plethora of proteins that have been implicated in EC signalling pathways, there may be value in examining EC RNAseq data to confirm that the genes for pathway-implicated proteins are indeed expressed in specific subtypes of endothelium, for example, in ECs of different organs. Whilst this would likely be a redundant exercise for proteins whose function has been defined within a range of different EC subtypes, this is a worthwhile endeavour for proteins whose EC roles are less clear or are controversial, as precedented by, for example, the debate surrounding the endogenous existence of anti-angiogenic VEGF isoforms (e.g. Bridgett et al., 2017) or when new candidates are discovered through genomic-wide association studies. However, the expression of some genes of interest may not be detected by RNAseq for technical reasons, such as insufficient read depth in bulk RNAseq assays (Hart et al., 2013, Toung et al., 2011) or gene drop out in single cell RNAseq assays (scRNAseq) (Marinov et al., 2014). Thus, multiple RNAseq resources must be examined to gain an accurate overview of the EC transcriptome (Khan et al., 2019).

Analysing the transcriptome of any given cell type is further complicated by the presence of low abundance transcripts that are proposed to be the product of leaky transcription (Hebenstreit et al., 2011, Hart et al., 2013, Gray et al., 2014). Unlike more moderately expressed genes, leaky genes are not associated with active chromatin markers and are not thought to be functional within the assayed cell type (Hebenstreit et al., 2011, Hart et al., 2013, Nagaraj et al., 2011). Instead, leaky transcription likely arises when an inactive gene resides near a highly expressed gene, with expression of the latter imparting a ‘transcriptional ripple effect’ on the former (Ebisuya et al., 2008, Hebenstreit et al., 2012, Gray et al., 2014). Several computational methods have been proposed for identifying leaky transcripts in bulk RNAseq data (Hebenstreit et al., 2011, Hart et al., 2013, Thompson et al., 2020, George and Chang, 2014). However, to the best of our knowledge, no study has systematically applied these methods to an exhaustive collection of bulk RNAseq for one cell type, nor has it been examined whether this approach could be used to systematically distinguish actively expressed from leaky genes in ECs.

Here, we have generated an exhaustive compendium of bulk RNAseq data sets from 5 endothelial subtypes and applied published methods (Hebenstreit et al., 2011, Hart et al., 2013) to classify EC genes as i. not expressed, ii. likely leaky expressed, or iii. likely actively expressed. We have termed this resource the <u>Bulk</u> RNAseq <u>E</u>ndothelial <u>C</u>ell explorer (BulkECexplorer). This resource will provide a convenient, fast and reliable method for evaluating whether genes of interest are actively expressed in ECs or EC subsets, in order to help prioritise genes for further investigation. Further, this resource provides a blueprint for developing analogous tools for other cell types.

## Results

Bulk RNAseq provides an average measure of gene expression in the cells of a given cell population and is thought to be well suited for detecting lowly expressed genes in that cell population (Marinov et al., 2014). However, an expressed gene may not be detected in any given bulk RNAseq for technical reasons. We therefore assembled an exhaustive compilation of publicly available bulk RNAseq data to generate a more accurate overview of gene expression for 5 EC subtypes commonly used for vascular biology research, including human dermal microvascular ECs (HDMECs), human umbilical vein ECs (HUVECs), primary mouse lung ECs, primary mouse brain ECs and primary mouse retina ECs.

To compile publicly available bulk RNAseq data from these EC subtypes into a compendium, we first queried the European Nucleotide Archive (ENA) for bulk RNAseq experiments that had analysed these EC subtypes. Our search returned 195 sequencing projects, with each project containing multiple RNAseq runs. After imposing our exclusion criteria (see methods), a total of 264 RNAseq runs with unique sample IDs were downloaded and aligned to the human or mouse genome, as appropriate. After quantifying transcript abundance, a further 24 samples were excluded from further analysis, because they lacked, or had very low levels of, transcripts for the core endothelial markers *KDR* or *CDH5*, or comprised a very low read number, or had poor read alignment. Thus, we selected a total of 240 endothelial bulk RNAseq samples from 59 sequencing projects for further analyses and compiled them into a compendium that is searchable via an online application, which we have termed the BulkECexplorer.

For each queried gene, the BulkECexplorer graphically displays for each EC subtype how many datasets contain the transcript and displays the transcript’s expression range in transcripts per million (TPM) (**Figure 1**). Further, the BulkECexplorer applies two established models to each dataset in the compendium to predict whether a queried gene is likely expressed at active or leaky levels in ECs (see below, **Figures 2** and **3**). All data are available for downloading in graph form or as a summary table.

**Figure 1.**
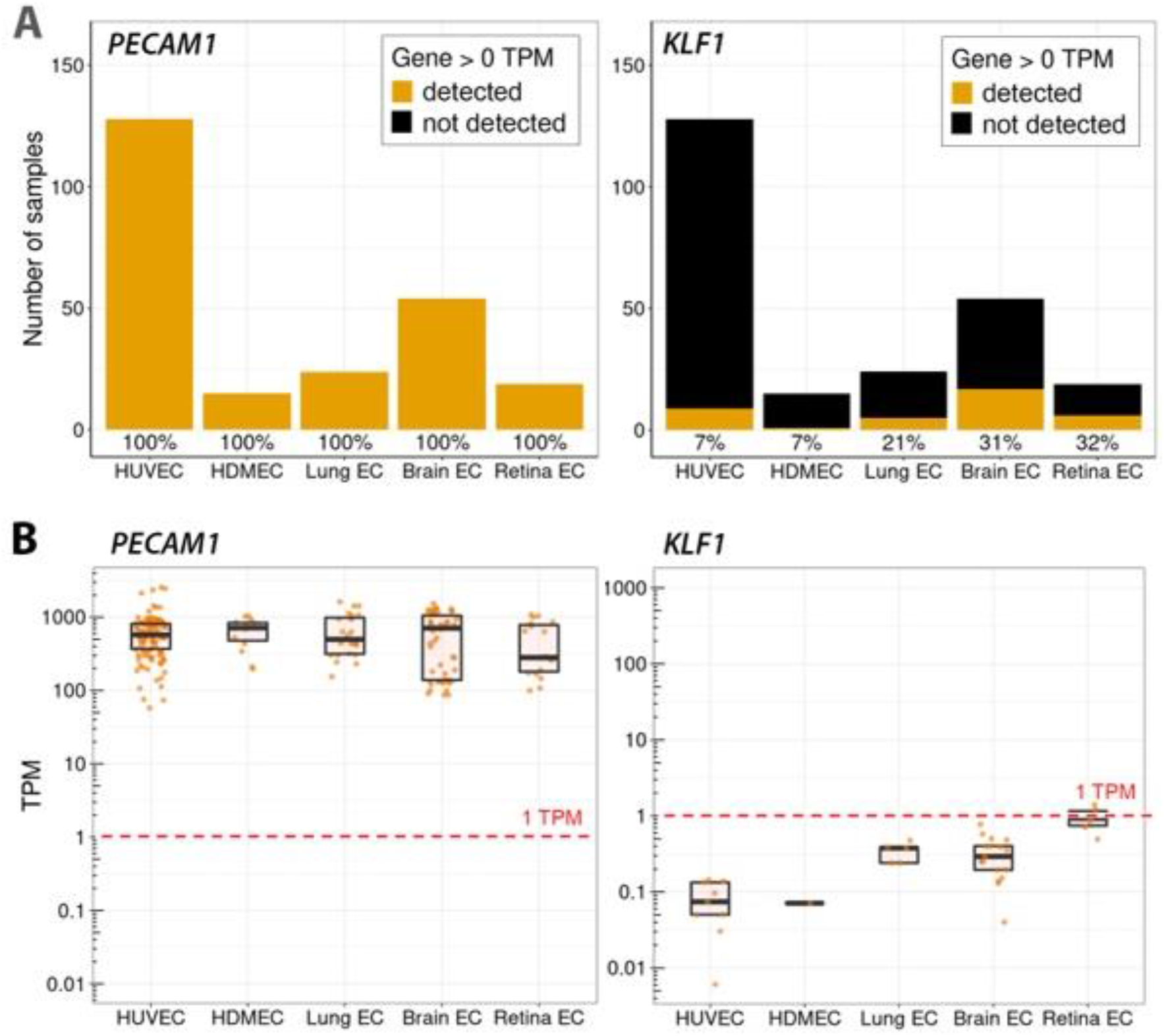
*PECAM1* and *KLF1* transcripts are detected in bulk RNAseq data from primary human and mouse ECs. Expression of *PECAM1* and *KLF1* in endothelial bulk RNAseq samples (HUVEC n = 128, HDMEC n = 15, lung EC = 24, brain EC = 54, retina EC n = 19). **(A)** Stacked bar charts depict the total number of samples per EC type and the frequency that each gene was detected or not detected in this EC type > 0 TPM (across EC subtypes; *PECAM1* n = 240, *KLF1* n = 38). **(B)** Expression values (TPM) for each gene in each sample for the indicated EC type, including boxplots to illustrate the interquartile range. The dashed red line indicates the 1 TPM threshold, a commonly used albeit heuristic transcript level to consider a gene to be expressed.

**Figure 2.**
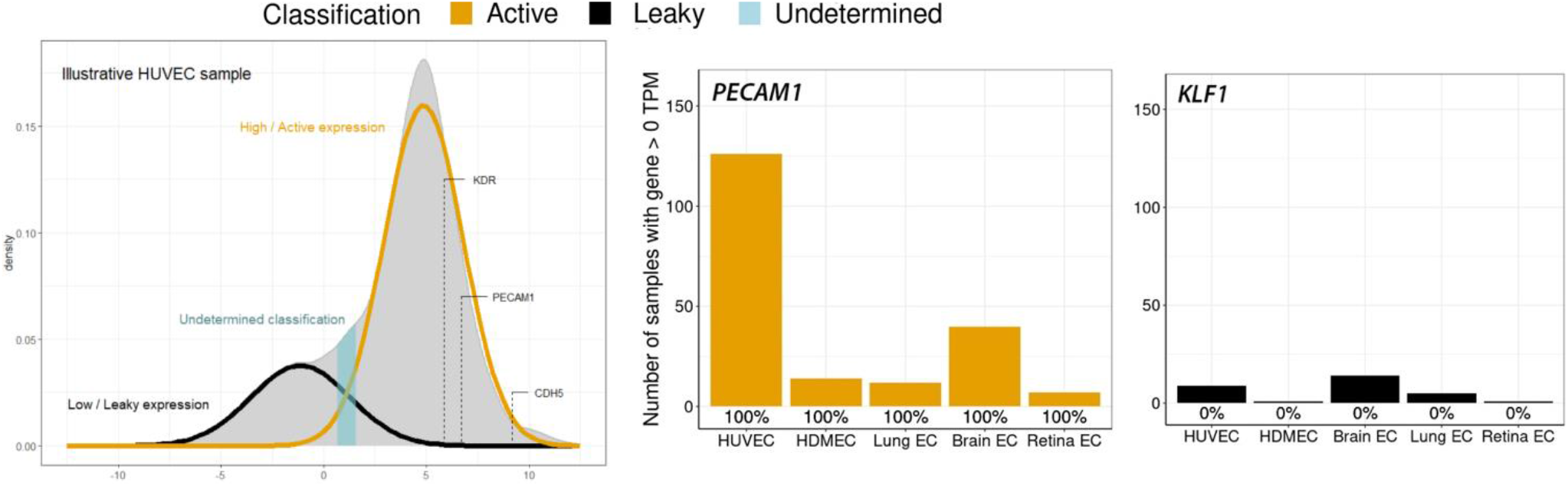
GMM-based classification suggests that *PECAM1* but not *KLF1* is actively expressed in ECs. Left: Kernal density estimates of the indicated log_2_-transformed bulk RNA seq samples. The expectation maximisation algorithm was used to estimate the parameters of the low and high Gaussian distributions (predicted leaky versus active expression), shown with black and gold lines, respectively. The log_2_TPM values for *PECAM1, KDR* and *CDH5* in each sample are indicated. Right: Stacked bar charts depicting the number of bimodally distributed samples per EC type in which the gene was classified as active, leaky or undetermined; the percentage of samples in which a gene was actively expressed in an EC type is reported below each bar (*PECAM1* n = 198, *KLF1* n = 30).

**Figure 3.**
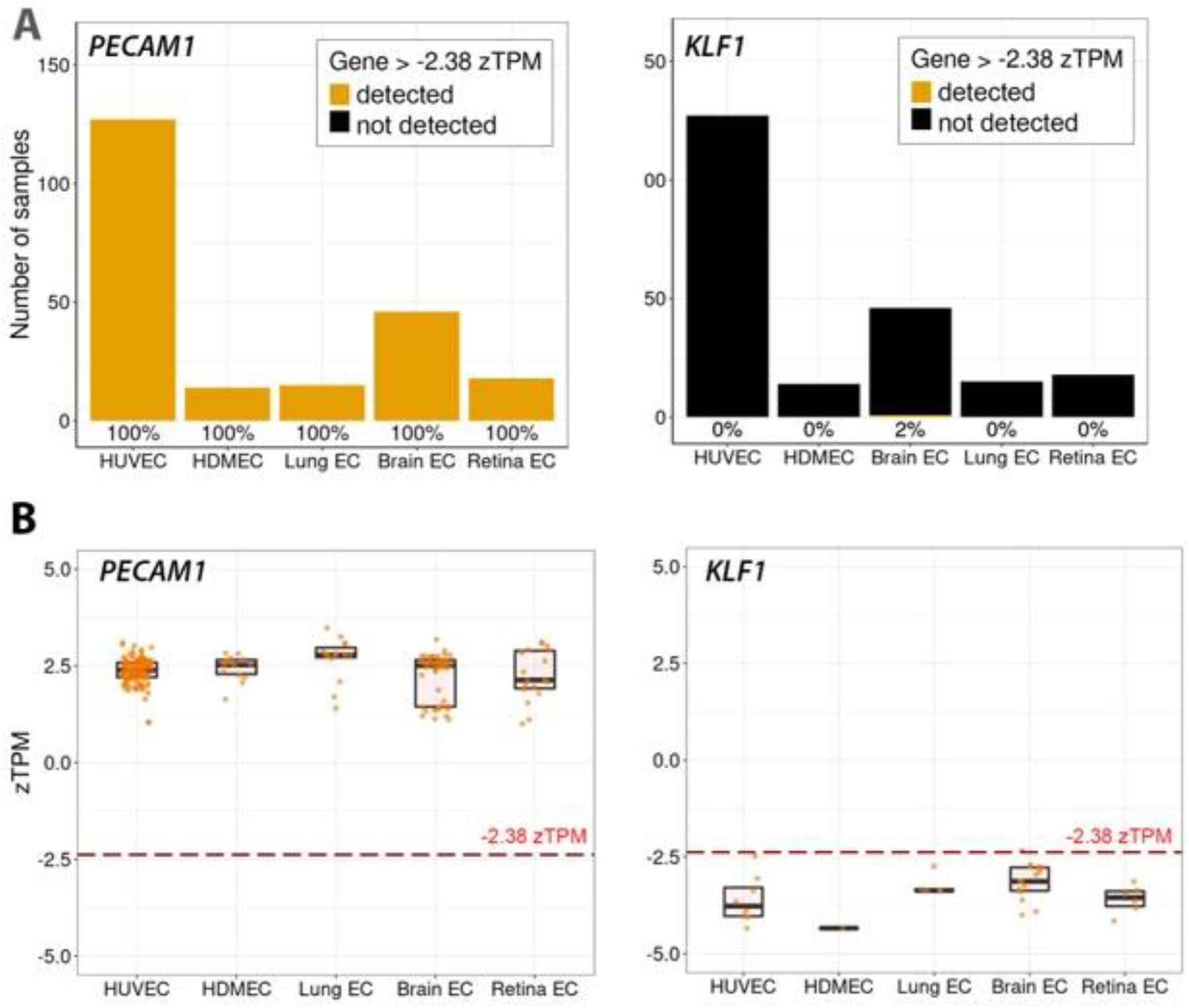
zTPM standardisation corroborates that *PECAM1* but not *KLF1* is actively expressed in ECs. **(A)** Stacked bar charts depicting the total number of samples per EC type in which each indicated gene was detected or not detected above the previously determined active expression threshold of -2.38 zTPM for ECs. The percentage of samples in which the indicated gene was actively expressed in an EC type is reported below each bar. **(B)** zTPM values for the indicated genes in each sample, split by EC type; values are shown as individual data points together with boxplots to illustrate the interquartile range. The dashed red line indicates the -2.38 zTPM threshold. Number of samples analysed across EC subtypes for each of the two genes n = 220 (datasets with a unimodal distribution were excluded); each data point represents one dataset.

The BulkECexplorer robustly detected *PECAM1* transcripts in all five EC subtypes (**Figure 1**). Instead, transcripts for the erythroid marker *KLF1* were detected in 15.8% of bulk RNA-seq datasets in the BulkECexplorer (**Figure 1A**; data resolved per EC type) at low levels (< 1 TPM), except in 2 mouse retina datasets, which had levels between 1 - 1.5 TPM (**Figure 1B**). Our *in silico* analysis with the BulkECexplorer compendium using detection > 0 TPM as a measure of gene expression could not reliably predict whether or not low transcript levels are indicative of leaky transcription of non-functional genes. To determine whether detected genes fit the profile of an actively or leaky transcribed gene, we applied published classification methods to the bulk RNAseq datasets in the BulkECexplorer. This approach is based on prior observations that the mixture of transcripts from leaky and actively expressed genes produces a bimodal distribution of transcript abundance in a homogenous population of mammalian cells (Hebenstreit et al., 2011). We first established that this bimodal distribution is also observed in the EC datasets included in the BulkECexplorer (**Figure 2** shows a representative HUVEC sample). Actively expressed genes are proposed to form a dominant Gaussian distribution in the higher expression range, whilst leaky genes are proposed to form a less prominent Gaussian distribution in the lower expression range; however, overlap between the two distributions produces a dominant right peak with a characteristic ‘left shoulder’ instead of two readily discernible distributions (Hebenstreit et al., 2011). Again, we established that this feature is observable in ECs (**Figure 2**). We can estimate the parameters of the leaky and active distributions by fitting a two component Gaussian mixture model (GMM) to the expression data, after which the probability of a gene belonging to either the active or leaky distributions can be calculated (Hebenstreit et al., 2011). We therefore applied this method to the bulk RNAseq datasets included in the BulkECexplorer (examples for the EC subtypes are shown in **Figure 2**).

We were able to fit a two component GMM to 98% of our compiled HDMEC and HUVEC samples and 61% of our compiled mouse EC samples (total: 199/240 samples). A further 23% of the mouse datasets presented a bimodal distribution with some degree of a left shoulder, but a third component was required to fit the GMM. As the nature of a third Gaussian distribution is undefined within the context of leaky versus active transcription, we did not include these samples in our analysis. The remaining mouse EC datasets appeared unimodal, without evidence of a left shoulder. As a unimodal distribution can reflect the presence of transcripts from a contaminating cell type (Hebenstreit et al., 2011, Hebenstreit and Teichmann, 2011), these datasets may reflect the technical pitfalls of separating a relatively small EC population from other, dominant cell types in small mouse organs. Thus, we restricted our analysis to the 198 human and mouse samples for which we could fit a two component GMM, indicative of leaky and active expression distributions. In each sample, a gene was classified as actively expressed if the probability of belonging to the high distribution, termed P(Active), was greater than 0.67, classified as leaky if P(Active) was less than 0.33, and classified as undetermined if P(Active) was between these two thresholds. Consistent with *PECAM1* expression being a defining criterion for EC identity, it was classified as actively expressed in all datasets of the BulkECexplorer (**Figure 2**; data resolved per EC type).

Demonstrating the reliability of the GMM method in assessing whether a lowly expressed gene is actively expressed in ECs or not, numerous genes not expected to be functional within ECs were correctly classified by the BulkECexplorer as leaky or non-expressed genes. Accordingly, the erythroid specific *KLF1* genes was classified as leaky in all the 38 datasets they were detected in (**Figure 2**; data resolved per EC type). Other examples of non-EC genes that were detected in some bulk RNAseq EC samples, but largely not classified as actively expressed when expressed at all, include another erythroid gene (*RHD*), ocular genes (*LENEP, CRYBB2*), osteoblast genes (*BGLAP, DMP1*) and several sex cell specific genes (*DDX4, GDF9, YBX2, SPACA4*) (see data in the BulkECexplorer). These findings support the validity of the GMM approach for classifying active vs. leaky gene expression.

The zFPKM algorithm provides an alternative method for distinguishing leaky versus actively expressed genes in bulk RNAseq data (Hart et al., 2013). With this algorithm, gene expression values are transformed into z-scores (zFPKM) based upon the parameters of the active expression Gaussian distribution, thereby providing a standardised measure of gene expression relative to the global pattern of gene expression in that sample (Hart et al., 2013). Using epigenomic and RNAseq data from the ENCODE project, a selection of cell-specific zFPKM thresholds have been calculated that reflect the value at which genes are more frequently associated with active rather than repressive chromatin, which is indicative of active versus leaky gene (Ernst et al., 2011, Consortium, 2012). For HUVECs, the only EC type analysed in the aforementioned study, the reported threshold was -2.38 zFPKM (Hart et al., 2013). Strong correlation between zTPM and zFPKM values (**Supplemental Figure 1**) indicates that the expression thresholds defined in zFPKM are applicable to zTPM values and allowed us to adopt the -2.38 threshold after applying the zTPM transformation to our compendium of EC bulk RNAseq datasets. Thereby, we found that *PECAM1* exceeded the -2.38 zTPM threshold in 100% of the samples in which it was detected (**Figure 3**; data resolved per EC type). By contrast, *KLF1* exceeded this threshold in only 0.5% of samples in which it was detected (**Figure 3**; data resolved per EC type).

In summary, we demonstrated that the BulkECexplorer is useful to compare the expression of genes between EC subtypes Further, we showed that the BulkECexplorer can predict whether gene expression is likely active, as expected from a functional gene within ECs, or leaky, consistent with an inactive gene in ECs.

## Discussion

Here, we have developed a compendium of publicly available bulk RNAseq data for 5 commonly studied EC subtypes. This resource can be interrogated as an *in silico* application to determine whether a gene is expressed in ECs, and whether detectable expression is likely to reflect active or leaky expression. We have termed this app the BulkECexplorer.

An interesting observation made when using the app was the low-level detection of non-EC genes detected within the datasets of the BulkECexplorer, including transcripts for the erythroid specific *KLF1*. Deep sequencing studies have hinted that most areas of the genome can be transcribed in a given cell type (Mercer et al., 2011), with some studies reporting >20,000 expressed genes in a homogenous cell population (Toung et al., 2011). Indeed, across our compilation of HUVEC samples alone, we detected transcripts for 19,436 genes out of a possible total of 19,878 protein-coding genes. Reports of expansive gene expression within a single cell type may appear difficult to reconcile with the concept of a cell-specific transcriptome, until it is appreciated that many of the unexpected transcripts are detected at very low levels. Thus, it has been suggested that two classes of protein-coding transcripts exist within a homogenous cell population – a higher expressed class that encode the functional proteome of the cell type, and a lower expressed class that is akin to biological noise, is likely non-functional and is caused by leaky transcription (Hebenstreit et al., 2011, Hart et al., 2013). Evidence to support a 2-class transcript model has been sourced from epigenomics and proteomics, where it has been reported that the lower expressed class of transcripts are not associated with epigenetic markers of active transcription and cannot readily be detected by mass spectrometry (Hebenstreit et al., 2011, Hart et al., 2013, Nagaraj et al., 2011, Sharma et al., 2015, Walley et al., 2016). Further, several studies have reported the presence of transcripts in the low expression cluster that are well characterized as specific markers of other cell types (Nagaraj et al., 2011, Hebenstreit et al., 2011). Indeed, we identified markers of various non-EC cell types in the low expression class of genes, including osteoblast, ocular and sex cell genes.

When analysing bulk RNAseq data, it can be predicted which class a gene likely belongs to by fitting a two-component GMM to the data, or by transforming the data using the zFPKM algorithm (Hart et al., 2013, Hebenstreit et al., 2011). Applying these methods to our EC bulk RNAseq compendium, the erythroid specific *KLF1* was consistently classified as non-expressed or a leaky gene in ECs. We conclude that the BulkECexplorer provides a reliable resource for comparing the distribution of specific genes amongst the EC subtypes that provide the most widely used models in vascular biology research to identify a suitable organ and in vitro model for further investigation. Further, the BulkECexplorer will complement ongoing scRNAseq studies to investigate whether candidate genes, predicted to be present in ECs but lacking from scRNAseq datasets are likely functional genes that remain undetected due to gene dropout for one of several conceivable reasons (Qiu, 2020). Another use of the BulkECexplorer would be to assess whether a candidate gene identified through a screen, such as genome wide association studies or 2-hybrid studies, is likely to be actively expressed in ECs and therefore warrants further investigation. Beyond the usefulness for the wider vascular community, this resource should provide a blueprint for developing analogous tools for other cell types.

## Methods

Bulk RNAseq datasets were retrieved from the European Nucleotide Archive (ENA) in July 2020. To identify relevant datasets, we queried the archive for the following terms: ‘HUVEC’, ‘HDMEC’, ‘HMVEC’, ‘dermal endothelial’, ‘retinal endothelial cells’, ‘brain endothelial cells’ or ‘mouse lung endothelial cells’. Our queries returned 195 RNAseq projects, whose datasets we individually examined to determine their suitability for our analysis. Only datasets generated by bulk or Ribo-tag RNAseq were retained for analysis. A small number of projects that contained samples with multiple run IDs were excluded to simplify and streamline downstream analysis. We also excluded datasets that were erroneously tagged as endothelial but did not include an EC type or which were ambiguous in their description. We only included murine datasets for brain, retina and lung ECs. As we wished to examine the ‘basal’ transcriptome of ECs, we excluded samples from rapidly growing and remodelling embryos. For the same reason, we excluded samples that had been stimulated (e.g., with a small molecule or by hypoxia) or had been genetically modified (e.g., by gene deletion, protein overexpression, siRNA-mediated knockdown or immortalisation). However, we retained samples in these projects that were derived from control cells (e.g., vehicle-stimulated or si-control transfected ECs). A total of 264 samples with a unique ID passed this exclusion stage. After alignment and transcript quantification, we further excluded samples that did not express >1 TPM of the core endothelial markers *KDR* or *CDH5*. We additionally excluded the study PRJEB14163, which contained samples with absent or low *KDR* expression and low read number. A total of 240 samples with unique ID from 59 projects passed this exclusion stage and were processed for further analysis (**Supplemental Table 1**). For this, FASTQ files were downloaded from the ENA. Reads were aligned to the human GRCh38 or mouse GRCm38 reference genomes, as appropriate, using HISAT2 version 2.1.0 (Kim et al., 2019). Transcript abundance was quantified using Stringtie version 2.1.3 (Pertea et al., 2015) with the reference annotation file Homo_sapiens.GRCh38.100.gtf or Mus_musculus.GRCm38.100.gtf, as appropriate (Ensemble). Transcript abundance was recorded as Transcripts per Million (TPM). RNAseq data were analysed in R Studio. To fit GMMs, expectation maximisation was performed on log_2_-transformed TPM values using the R package ‘Mixtools’ and the *normalmixEM* function set to find 2 components/distributions. Samples that displayed a unimodal log_2_TPM distribution or required fitting with >2 components/distributions were excluded from this analysis, because a low- and high-expression gene clusters could not be readily identified (n = 42 excluded, n = 198 analysed; Supplemental Table 2). zTPM scores were calculated using the R ‘zFPKM’ package (Hart et al., 2013). Samples that displayed a unimodal log_2_TPM distribution were excluded from this analysis, because the zTPM transformation relies on identifying a high expression Gaussian distribution (n = 20 excluded, n = 220 included; Supplemental Table 2). The R ‘shiny’ and ‘shinydashboard’ packages were used to create the BulkECexplorer web application.

## Acknowledgements

This study was supported by grants from the British Heart Foundation [PG/19/37/3439], Medical Research Council [MR/N011511/] and Wellcome [205099/Z/16/Z] to CR, the British Heart Foundation to CR and AF [PG/18/85/34127], the Fondazione Cariplo (2018-0298) and the Fondazione Associazione Italiana per la Ricerca sul Cancro (AIRC) (22905) to AF and a PhD fellowship to GDP [FS/4yPhD/F/21/34157].

## Authors’ contributions

JTB, AF and CR conceived and designed the study, analysed the data and co-wrote the manuscript. JTB and GDP wrote the code. All authors read and approved the submitted manuscript.

---

**Supplementary Figure 1.**
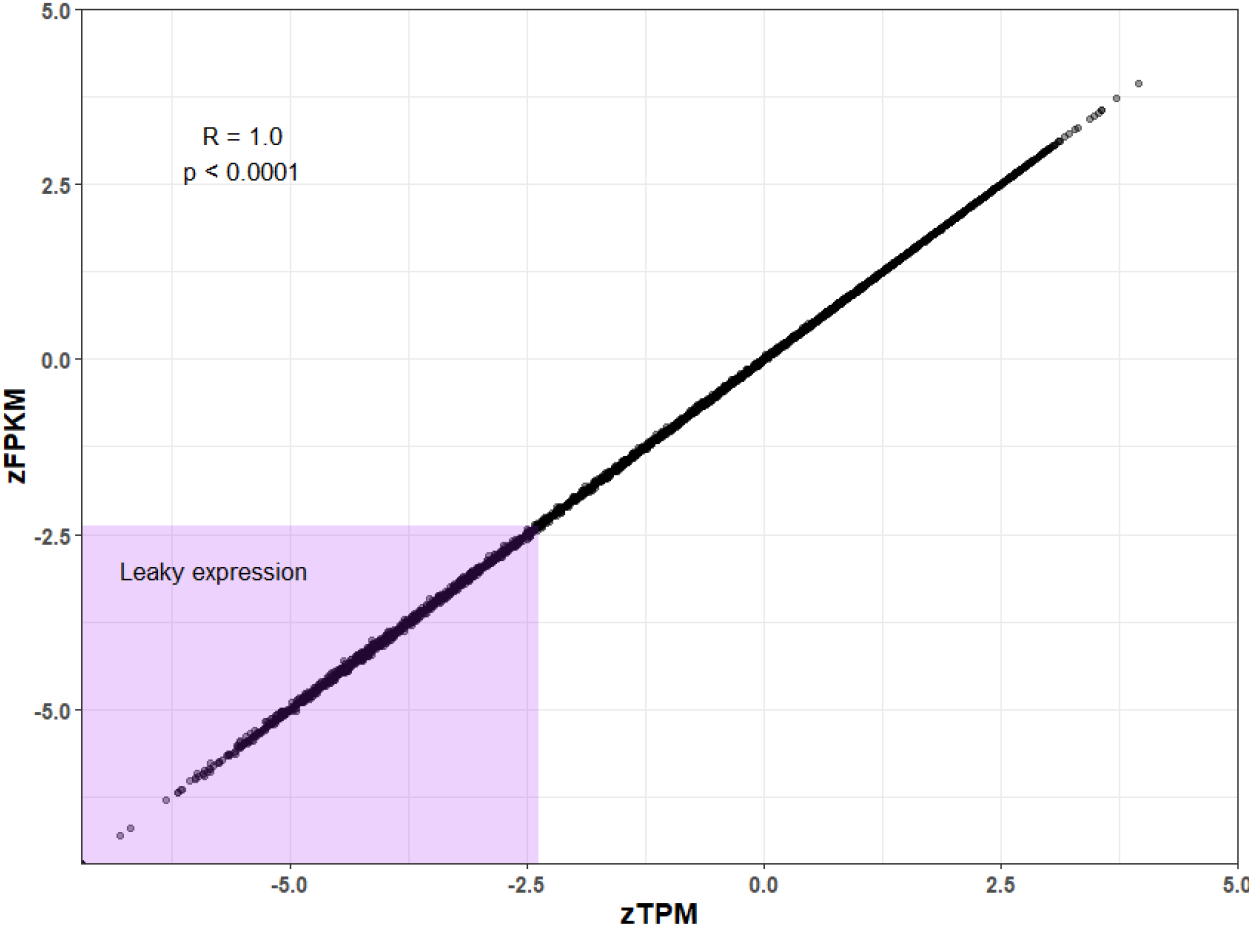
zTPM correlates with zFPKM. Correlation of zTPM and zFPKM units for a selection of 37 vascular, neuronal and immune genes in our compendium of bulk RNAseq datasets (n = 220, unimodal datasets not analysed); the Pearson correlation coefficient is provided (R). Each data point corresponds to a single gene within a single dataset. The purple box highlights values below the published threshold for leaky expression in HUVECs of -2.38.

